# The gatekeeper of *Yersinia* type III secretion is under RNA thermometer control

**DOI:** 10.1101/2021.05.20.444929

**Authors:** Stephan Pienkoß, Soheila Javadi, Paweena Chaoprasid, Thomas Nolte, Christian Twittenhoff, Petra Dersch, Franz Narberhaus

**Affiliations:** Microbial Biology, Ruhr University Bochum, Germany; Institute of Infectiology, Center for Molecular Biology of Inflammation (ZMBE), University of Münster, Münster, Germany; Rottendorf Pharma GmbH, Ennigerloh, Germany

## Abstract

Many bacterial pathogens use a type III secretion system (T3SS) as molecular syringe to inject effector proteins into the host cell. In the foodborne pathogen *Yersinia pseudotuberculosis,* delivery of the secreted effector protein cocktail through the T3SS depends on YopN, a molecular gatekeeper that controls access to the secretion channel from the bacterial cytoplasm. Here, we show that several checkpoints adjust *yopN* expression to virulence conditions. A dominant cue is the host body temperature. A temperature of 37 °C is known to induce the RNA thermometer (RNAT)-dependent synthesis of LcrF, a transcription factor that activates expression of the entire T3SS regulon. Here, we uncovered a second layer of temperature control. We show that another RNAT silences translation of the *yopN* mRNA at low environmental temperatures. The long and short 5’-untranslated region of both cellular *yopN* isoforms fold into a similar secondary structure that blocks ribosome binding. The hairpin structure with an internal loop melts at 37 °C and thereby permits formation of the translation initiation complex as shown by mutational analysis, *in vitro* structure probing and toeprinting methods. Importantly, we demonstrate the physiological relevance of the RNAT in the faithful control of type III secretion by using a point-mutated thermostable RNAT variant with a trapped SD sequence. Abrogated YopN production in this strain led to unrestricted effector protein secretion into the medium, bacterial growth arrest and delayed translocation into eukaryotic host cells. Cumulatively, our results show that substrate delivery by the *Yersinia* T3SS is under hierarchical surveillance of two RNATs.

**Author summary:** Temperature serves as reliable external cue for pathogenic bacteria to recognize the entry into or exit from a warm-blooded host. At the molecular level, a temperature of 37 °C induces various virulence-related processes that manipulate host cell physiology. Here, we demonstrate the temperature-dependent synthesis of the secretion regulator YopN in the foodborne pathogen *Yersinia pseudotuberculosis*, a close relative of *Yersinia pestis*. YopN blocks secretion of effector proteins through the type III secretion system unless host cell contact is established. Temperature-specific regulation relies on an RNA structure in the 5’-untranslated region of the *yopN* mRNA, referred to as RNA thermometer, which allows ribosome binding and thus translation initiation only at an infection-relevant temperature of 37 °C. A mutated variant of the thermosensor resulting in a closed conformation prevented synthesis of the molecular gatekeeper YopN and led to permanent secretion and defective translocation of virulence factors into host cells. We suggest that the RNA thermometer plays a critical role in adjusting the optimal cellular concentration of a surveillance factor that maintains the controlled translocation of virulence factors.

## Introduction

Numerous gram-negative plant-or animal pathogens deploy a type III secretion system (T3SS) to translocate effector proteins into eukaryotic host cells [1,2]. This molecular syringe or “injectisome” is evolutionary related to flagella. The complex nanomachines crosses the bacterial envelope and the host membrane for cross-kingdom transfer of effectors into the host cell cytosol [3]. While the overall architecture and the structural components of the T3SS are conserved, the cocktail of secreted proteins varies in order to execute diverse species-specific activities. Among other processes, effector proteins can target the host cytoskeleton, autophagy and innate immune response [4–7].

T3SS components in members of the genus *Yersinia* are encoded on the virulence plasmid [8]. This genus consists of the human pathogens *Yersina pestis,* the causative agent of bubonic and pneumonic plague [9], and of the enteropathogens *Yersinia enterocolitica* and *Yersinia pseudotuberculosis* [10]. The T3SS is composed of more than 15 different proteins and the biogenesis of this more than 15 MDa-large apparatus is a strictly regulated hierarchical process that is controlled by internal and external cues at various transcriptional and posttranscriptional levels [11,12]. The basal body is composed of more than a dozen so-called Ysc proteins and spans the bacterial inner and outer membranes. It is followed by the needle structure, a polymer of the YscF protein. The distal end of the needle serves as platform for the pore complex comprised of the LcrV-containing tip and YopBD, the translocation pore [13]. The formation of the needle and the pore complex as well as the secretion of effector proteins follow a specific secretion order. First, substrates consisting of the needle subunit YscF and the ruler protein YscP, which determines the length of the needle, are secreted [3,14–16]. Once the contact with the host cell is established, proteins of the pore complex pass through the T3SS and integrate into the host membrane to allow the passage of effector proteins [17–19].

Yet another checkpoint regulates the fidelity of type III secretion (T3S) and prevents the release of *Yersinia* outer proteins (Yops) prior to host cell contact (Fig. 1). YopN forms a heterodimer with TyeA that together with the SycN/YscB chaperones plugs the secretion channel until secretion is desired [20,21]. While TyeA binds to the C-terminus of YopN, the chaperone complex SycN/YscB binds to the N-terminus ensuring attachment to the T3SS [21–23]. Once contact with the host cell is made, YopN-TyeA dissociates and SycN/YscB facilitates the export of YopN through the T3SS. Translational +1 frameshifting near the 3’ end of the *yopN* mRNA in *Y. pestis* and *Y. pseudotuberculosis* occasionally generates a YopN-TyeA hybrid protein that maintains secretion control suggesting that the reversible interaction between YopN and TyeA is not a functional prerequisite [24,25].

**Fig 1.**
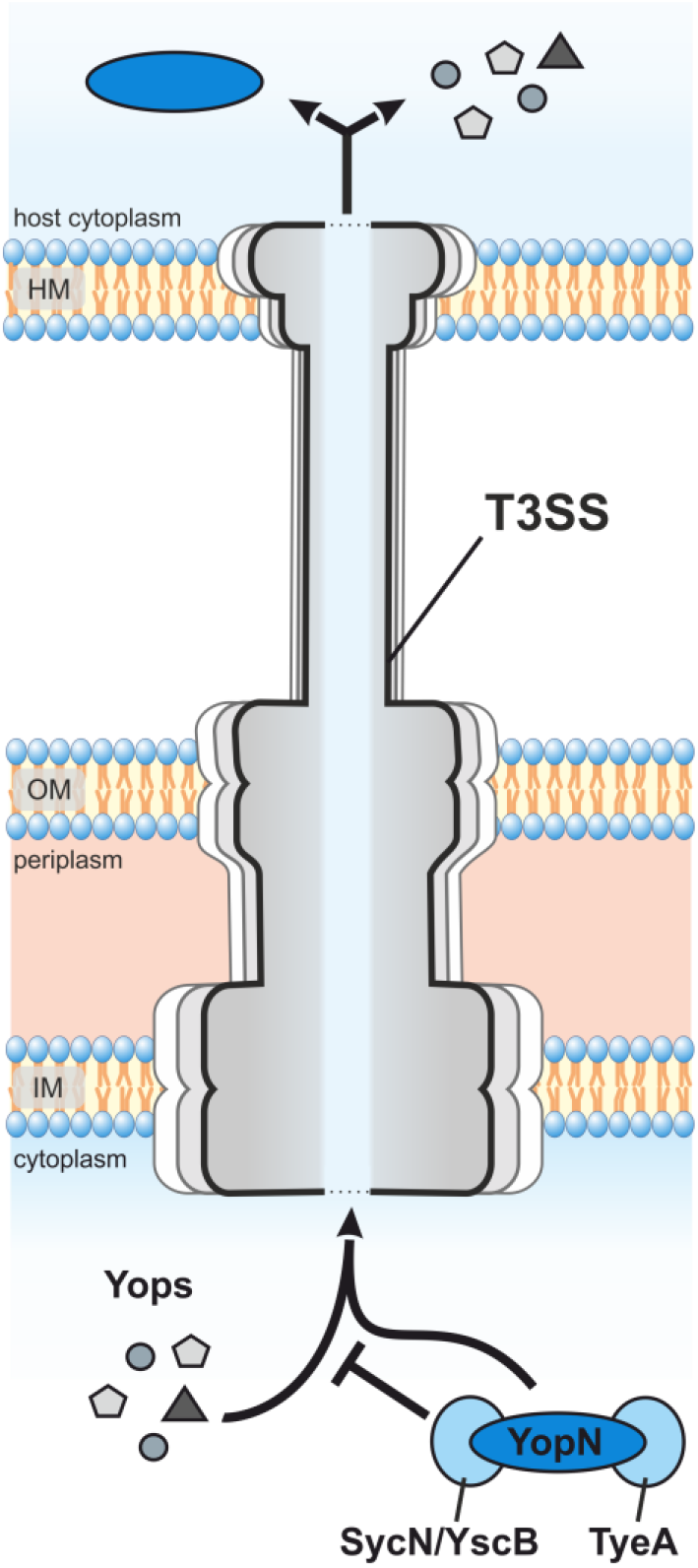
Schematic overview of YopN-regulated Yop secretion in *Yersinia.* The secretion of effector proteins (Yops) that modulate the host immune system are controlled by the YopN-TyeA-SycN-YscB complex. YopN and TyeA block secretion in concert with the chaperone complex SycN/YscB [20,22]. Dissociation of the plug is assumed to occur through depletion of calcium and after host cell contact resulting in the SycN/YscB-controlled secretion of YopN itself [23,26,27]. HM: host membrane; OM: outer membrane; IM: inner membrane.

Virulence conditions can be mimicked in the laboratory by decreasing the calcium concentration in the surrounding medium, which is why the YopN complex has been described as calcium plug [26,27]. YopN export subsequently enables the secretion of other Yops [22,28]. Deletion of *yopN* results in a temperature-sensitive phenotype at 37 °C characterized by continuous secretion of effector proteins associated with growth impairment [27]. YopN might serve additional functions within the host cell. Recently, it was shown to affect systemic infection in mice and to adjust the secretion of the effector proteins YopE and YopH [29,30].

Temperature is a major stimulus for *Yersinia* T3SS gene expression both under *in vitro* laboratory conditions and in the host [31–33]. The pathogen interprets a temperature of 37 °C as signal that it has been taken up by a warm-blooded host. Temperature-dependent transcription initiation of a large array of virulence genes is accomplished by the AraC-type regulator LcrF, whose expression is under the influence of an RNA thermometer (RNAT) [34]. RNATs are thermolabile RNA structures in the 5’-untranslated region (5’-UTR) that sequester the Shine Dalgarno (SD) sequence and/or the start codon. At ambient environmental temperatures, ribosome binding to the SD sequence is prevented by the double-stranded RNA. The metastable structure melts in the host at about 37 °C, which liberates the ribosome binding site and permits translation initiation [35,36].

In search of new temperature-responsive RNA structures in *Y. pseudotuberculosis,* we used two global RNA structuromics approaches and identified numerous RNAT candidates that undergo a conformational change from 25 to 37 °C [37,38]. Several of them are located upstream of genes critical for various aspects of bacterial virulence. A recently documented example is the *cnfY* thermometer controlling translation of the cytotoxic necrotizing factor that enhances inflammation and Yop delivery by activation of Rho GTPases in the host [39–41]. In contrast to Yops, the CnfY toxin is not secreted by the T3SS but delivered to host cells via outer membrane vesicles [42].

Another promising RNAT candidate was identified in the 5’-UTR of *yopN* coding for the T3SS regulator described above. In this study, we show the functionality of this RNAT in both naturally occurring short and long transcripts of *yopN* by various *in vitro* and *in vivo* assays at different temperatures. Most importantly, by using thermostable RNAT variants with a sequestered SD sequence, we demonstrate the physiological relevance of the RNAT in the proper control of type III secretion.

## Results

### A putative RNAT in the short and long 5’-UTR of *yopN*

*Y. pseudotuberculosis yopN* is the first gene of the heptacistronic *virA* operon composed of *yopN, tyeA, sycN, yscX, yscY, yscV* and *lcrR* [43]. Two alternative transcription start sites upstream of *yopN* result in a short and a long 5’-UTR of 37 and 102 nucleotides, respectively (Fig. 2A). Previous RNA-Seq analyses showed that the short transcript is more highly expressed at 37 °C than the long transcript [32,37]. We examined the expression of both *yopN* isoforms by qRT-PCR and, consistent with the previous studies, saw an over-abundance of the short transcript (Fig. 2B). Both transcripts were barely detectable at 25 °C. The amount of the short isoform increased about 22-fold at 37 °C under non-secretion conditions (+ Ca^2+^) and even further to 63-fold under secretion-mimicking conditions (-Ca^2+^). The long transcript was poorly transcribed and three-fold induced at 37 °C in the presence of Ca^2+^. Overall, these results show that the short transcript is the major isoform responsible of *yopN* expression. A striking observation of our previous comparative analysis of the *Y. pseudotuberculosis* RNA structurome at 25 and 37 °C [37] was a thermolabile RNA element upstream of *yopN.* The SD sequence (5’-AGGGAGU-3’) and the start codon are part of the same hairpin structure in both the short and the long transcript (Fig. 2C and 2D). An internal loop that exposes some nucleotides of the SD sequence might explain the temperature sensitivity of this structure. Compared to the short transcript, the longer one features a slightly extended hairpin, two additional hairpins downstream of it and a base-paired region between the immediate 5’ end and nucleotides in the early coding region of *yopN.* A sequence comparison showed that the *yopN* 5’-UTRs of *Y. pseudotuberculosis* and *Y. pestis* are identical over the entire length of the long transcript and at least 30 nts into the coding region (Fig. S1). Three mutations were found in the corresponding *Y. enterocolitica* sequence. In particular, the exchange of residue U6 opposite the first nucleotide of the AUG start codon for a C residue might reduce the overall stability of the RNA structure (Fig. 2C).

**Fig 2.**
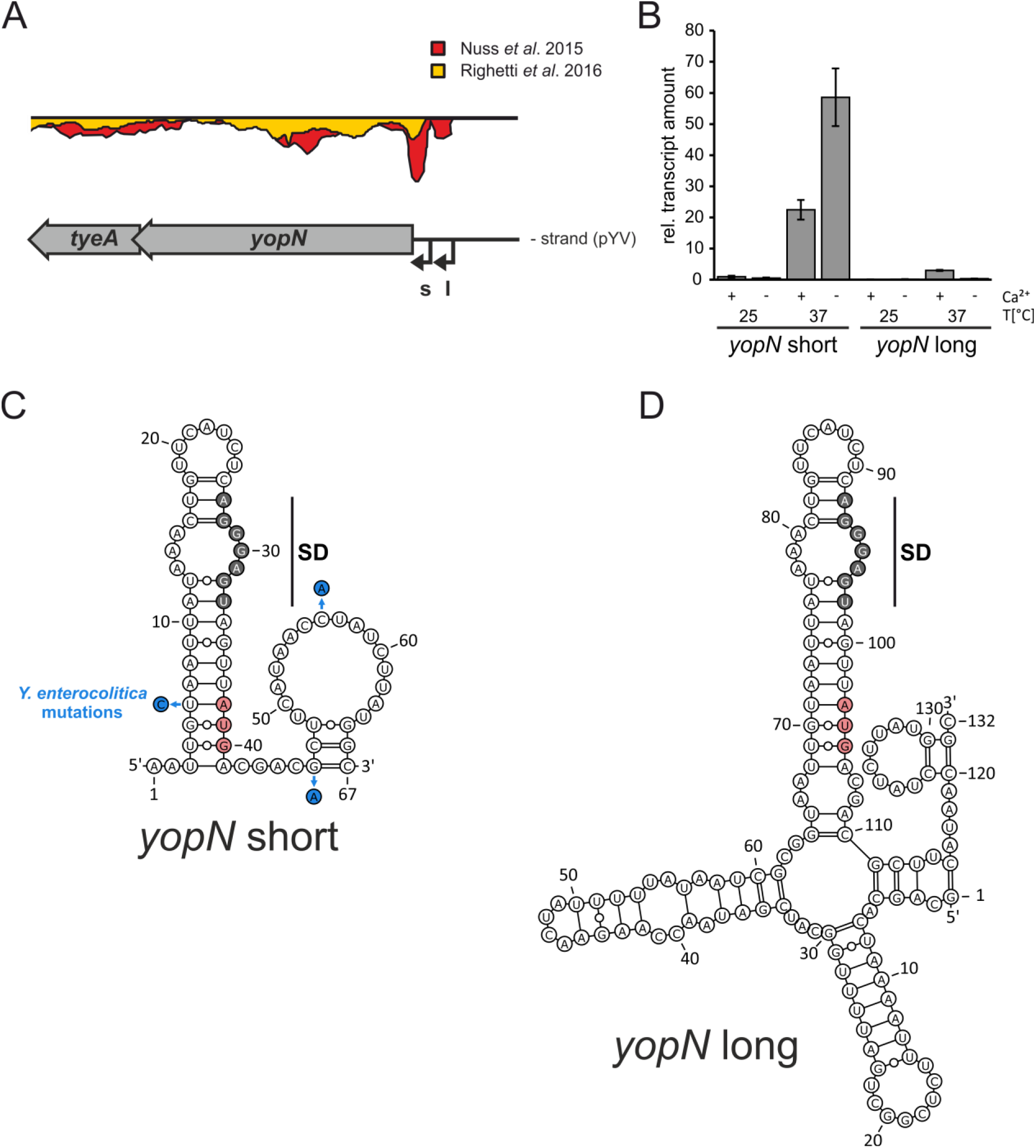
Expression and RNA structures of 5’-UTRs of *yopN.* (A) RNA-seq results of the *yopN* locus at 37 °C and identification of two transcriptional start sites from [32] and [37] visualized by the Artemis genome browser (s: short transcript; l: long transcript). (B) Comparison of the relative transcript levels of both *yopN* 5’-UTRs under non-secretion (+ Ca^2+^) and secretion (- Ca^2+^) conditions at 25 °C and 37 °C. Samples of *Y. pseudotuberculosis* YPIII were taken during the early exponential phase at an OD_600_ of 0.5 followed by RNA isolation and qRT-PCR. Transcript levels were normalized to the amount of *yopN* short at 25 °C under non-secretion conditions and to the reference genes *nuoB* and *gyrB.* The mean transcript amounts and standard deviations comprise the results of three biological replicates. (C,D) PARS-derived RNA secondary structures of the short and long 5’-UTRs of *yopN* at 37 °C including the first 30 nucleotides of the coding region [37]. The putative SD region is highlighted in gray and the start codon in red. Nucleotide exchanges in the *Y. enterocolitica* sequence are indicated in (C).

### Predictable consequences of point mutations in the *yopN* RNAT of both transcripts

The structured 5’-UTRs of the short and long *yopN* transcripts suggest that the transcriptional control of *yopN* expression in response to temperature (Fig. 2A and B) is complemented by RNAT-mediated translational control. To investigate the functionality of the putative *yopN* RNAT, five variants of the hairpin of the short transcript were generated by site-directed mutagenesis (Fig. 3A). Three variants were designed to result in a putative stable hairpin (referred to as repressed; R1, R2 and R3). In R1 and R2, the internal loop of the hairpin is reduced (R2) or completely closed (R1) by residues pairing with the SD sequence. In R3, base pairing with the AUG start codon is strengthened. Two other variants were expected to result in a less stable hairpin (referred to as derepressed; D1 and D2). They result in a larger internal loop of the hairpin and partially (but not completely) liberate the SD sequence. The functionality of these variants was measured quantitatively by reporter gene assays using translational fusion constructs consisting of the *yopN* 5’-UTR (WT), the different *yopN* variants and the positive control *lcrF* fused to *bgaB* coding for a heat-stable β-galactosidase (Fig. 3B). Compared with the positive control *lcrF* that exhibited a temperature-dependent 3-fold induction of reporter enzyme activity, the *yopN* RNAT showed a 6.9-fold increased β-galactosidase activity at 37 °C compared to 25 °C (Fig 3C). The point-mutated variants behaved as expected. R1 with the completely paired SD sequence fully repressed expression at both temperatures whereas R2 that contained just one additional base pair in the internal loop reduced expression at 25 °C but remained inducible at 37 °C. Stabilizing the AUG start codon in R3 almost completely blocked expression. In contrast, variants D1 and D2 already showed elevated β-galactosidase activities at 25 °C, which further increased at 37 °C. The reporter gene activities were fully reflected in Western blot experiments detecting the His-tagged BgaB enzyme (Fig. 3C). Cumulatively, these results provide support for the existence of a functional RNAT in the short 5’-UTR of *yopN*.

**Fig 3.**
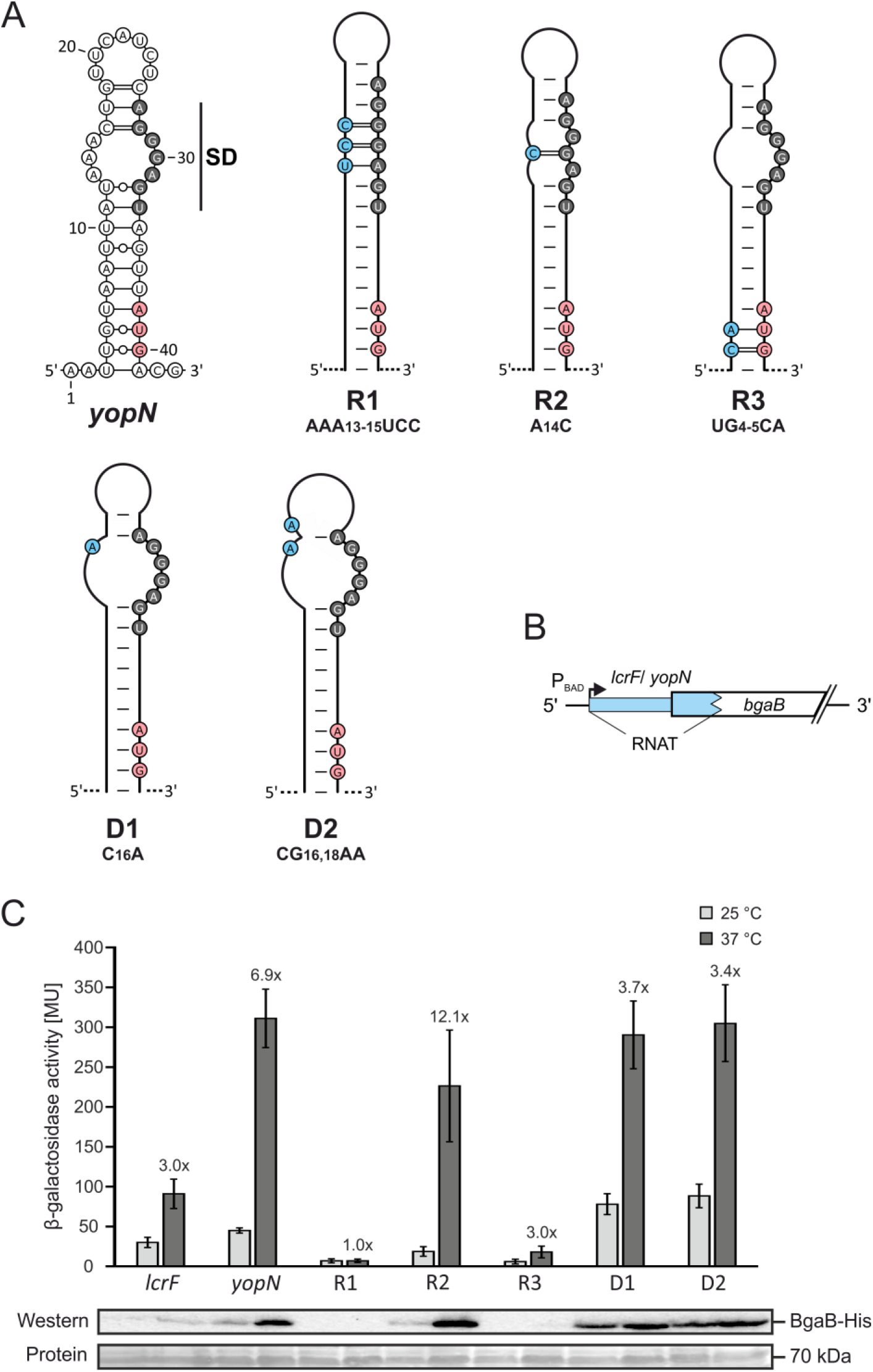
Schematic representation and functional characterization of the short *yopN* 5’-UTR and mutated variants. (A) PARS-derived stem-loop RNA structure of the short *yopN* 5’-UTR (nucleotides 1 to 43) [37] and predicted stabilized (R1 – 3) and destabilized (D1 – 2) variants. The putative SD region is highlighted in gray, the start codon in red and mutated nucleotides are highlighted in blue. (B) Plasmid-based translational fusions of 5’-UTRs of interest and *bgaB* encoding a heat-stable β-galactosidase to test RNA thermometer (RNAT) functionality. Expression of the fusion products is controlled by the arabinose-inducible promotor P_BAD_. The RNAT of *lcrF* served as a positive control [34]. (C) The β-galactosidase assays of *lcrF,* the short *yopN* 5’-UTR and the corresponding mutated variants were conducted at 25 and 37 °C. *Y. pseudotuberculosis* YPIII cells carrying plasmids of the fusion constructs were grown to an OD_600_ of 0.5 at 25 °C. Subsequently, transcription of the reporter gene was induced by 0.1 % (w/v) L-arabinose and the cultures were split to flasks at 25 and prewarmed flasks at 37 °C and incubated for further 30 minutes. Samples were then taken for the β-galactosidase assay. The mean activities in Miller Units and the mean standard deviations were calculated from nine biological replicates. The representative Western blot displays the amount of BgaB-His produced. Protein amounts were adjusted to an optical density of 0.5 and detected by Ponceau S staining after blotting onto a nitrocellulose membrane.

We then wondered whether the structure in the long *yopN* transcript (Fig. 2D) is equally temperature responsive and constructed two corresponding *bgaB* fusions to the WT region and a repressed R1 variant. The β-galactosidase experiments showed essentially the same results for the short and long 5’-UTRs (Fig. 4) strongly suggesting that they both contain functional RNATs.

**Fig 4.**
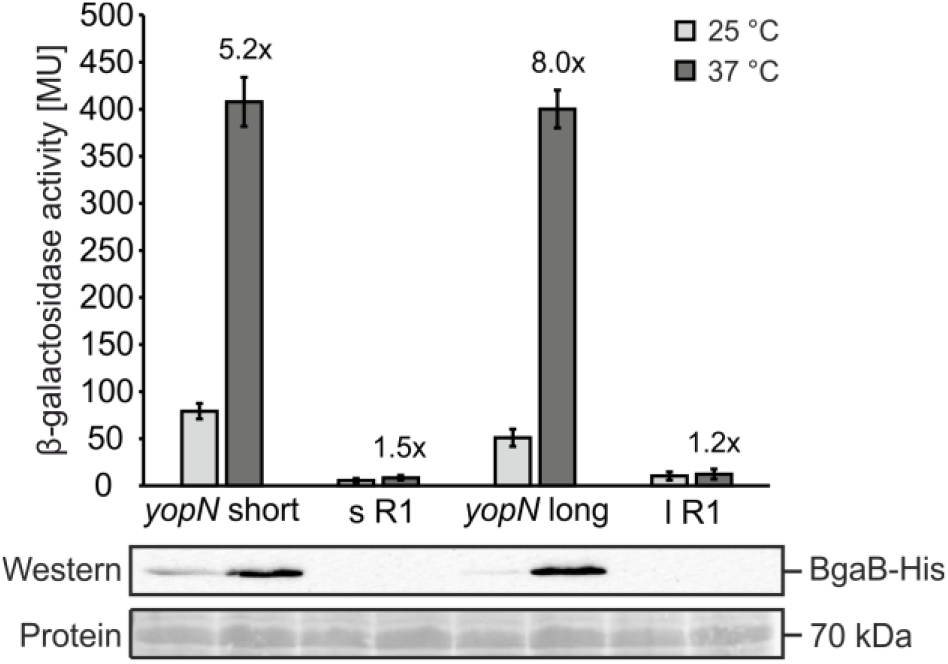
RNAT functionality of the short and long 5’-UTR of *yopN.* Translational *bgaB* fusions of the wild type and stable R1 UTRs were tested as described in the legend of Fig 3.

Finally, to support translational control and exclude any transcriptional control by the 5’-UTR, the short *yopN* RNAT and the R1 and D2 variants were translationally fused to *gfp* (Fig. 5A) and mRNA and protein levels were recorded (Fig. 5B). Importantly, when transcription was induced from the P_BAD_ promoter by arabinose addition, the transcript amounts of each construct were the same at 25 and 37 °C excluding an influence of temperature on transcription and/or transcript stability. In accordance with the BgaB results, GFP production occurred as predicted, i.e. inducible GFP levels at 37 °C in the WT situation, overall elevated GFP protein in the D2 strain and fully repressed GFP production in R1. All these results are in favour of a model, in which the thermolabile 5’-UTR upstream of *yopN* melts at host body temperature and thereby facilitates translation initiation.

**Fig 5.**
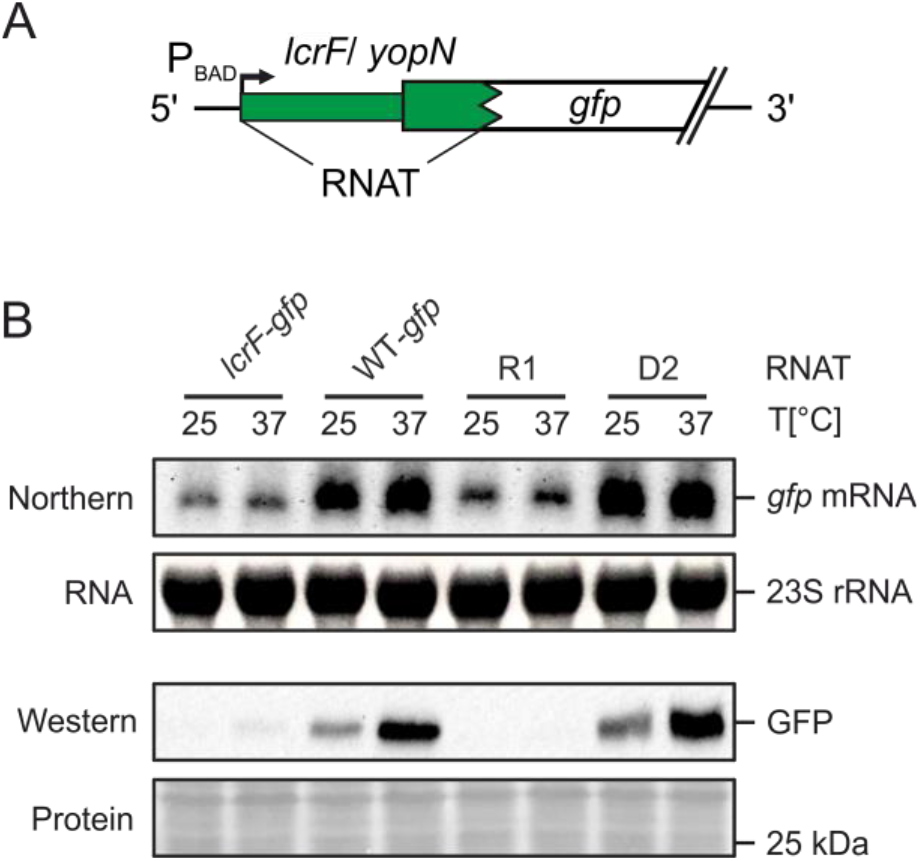
Temperature-controlled expression of *gfp* by the short *yopN* RNAT. (A) Plasmid-based translational fusions of *yopN:gfp* were cloned to test RNAT functionality at RNA and protein levels. Transcription of the fusion product is controlled by the arabinose inducible promotor P_BAD_. (B) Determination of transcript and protein levels of *yopN:gfp (WT-gfp)* and mutated *yopN* variants (R1 and D1) by Northern and Western blot analyses, respectively. The RNAT of *lcrF*served as a positive control [8]. *Y. pseudotuberculosis* YPIII cells carrying plasmids of the fusion constructs were grown to an OD_600_ of 0.5 at 25 °C. Transcription was induced by 0.1 % (w/v) L-arabinose and the cultures were split to flasks at 25 °C and prewarmed flasks at 37 °C and incubated for further 30 minutes. Samples were then taken for Northern and Western blot analyses. The blots shown represent one of three biological replicates. To ensure equal amounts of RNA, a total of 10 μg of RNA was loaded per sample. Ethidium bromide stained 23S rRNA served as loading control. Protein amounts were adjusted to an optical density of 0.5 and detected by Ponceau S staining after blotting onto a nitrocellulose membrane.

### Melting of the RNA structure liberates the SD sequence and start codon

To lend biochemical support to the gradual melting of the RNAT with elevated temperatures, we conducted enzymatic structure probing experiments on *in vitro*-synthesized RNAs (Fig. 6). The short WT and R1 RNATs were treated with RNases T1 and T2 at 25, 37 and 42 °C. RNase T1 specifically introduces cuts in single-stranded RNA (ssRNA) at the 3’ end of guanines, whereas T2 prefers ssRNA at the 3’ end of adenines but also cleaves other single-stranded nucleotides. The presence of cuts at all temperatures in the loop at the top of the hairpin and in the large loop at the beginning of the coding region (positions 19-25 and 49-64, respectively) suggested folding of the RNA into the anticipated structure (Fig. 6A and B). Poor cleavage at 25 °C and gradual heat-induced melting was observed in the region of the SD sequence at nucleotides 27-32 (5’-AGGGAG-3’) and in the corresponding anti-SD region with several A residues around position 14 (Fig. 6A and B; quantification for some of the bands in Fig. 6C). In addition, temperature-modulated cleavage was also observed for nucleotides of the stem structure at the bottom of the hairpin (position 34-40), which includes the AUG start codon. The start codon showed the same behavior in the R1 construct. The SD sequence in this RNA structure, however, was inaccessible to RNases even at high temperatures due to the introduced stabilizing point mutations (Fig. 3A).

**Fig 6.**
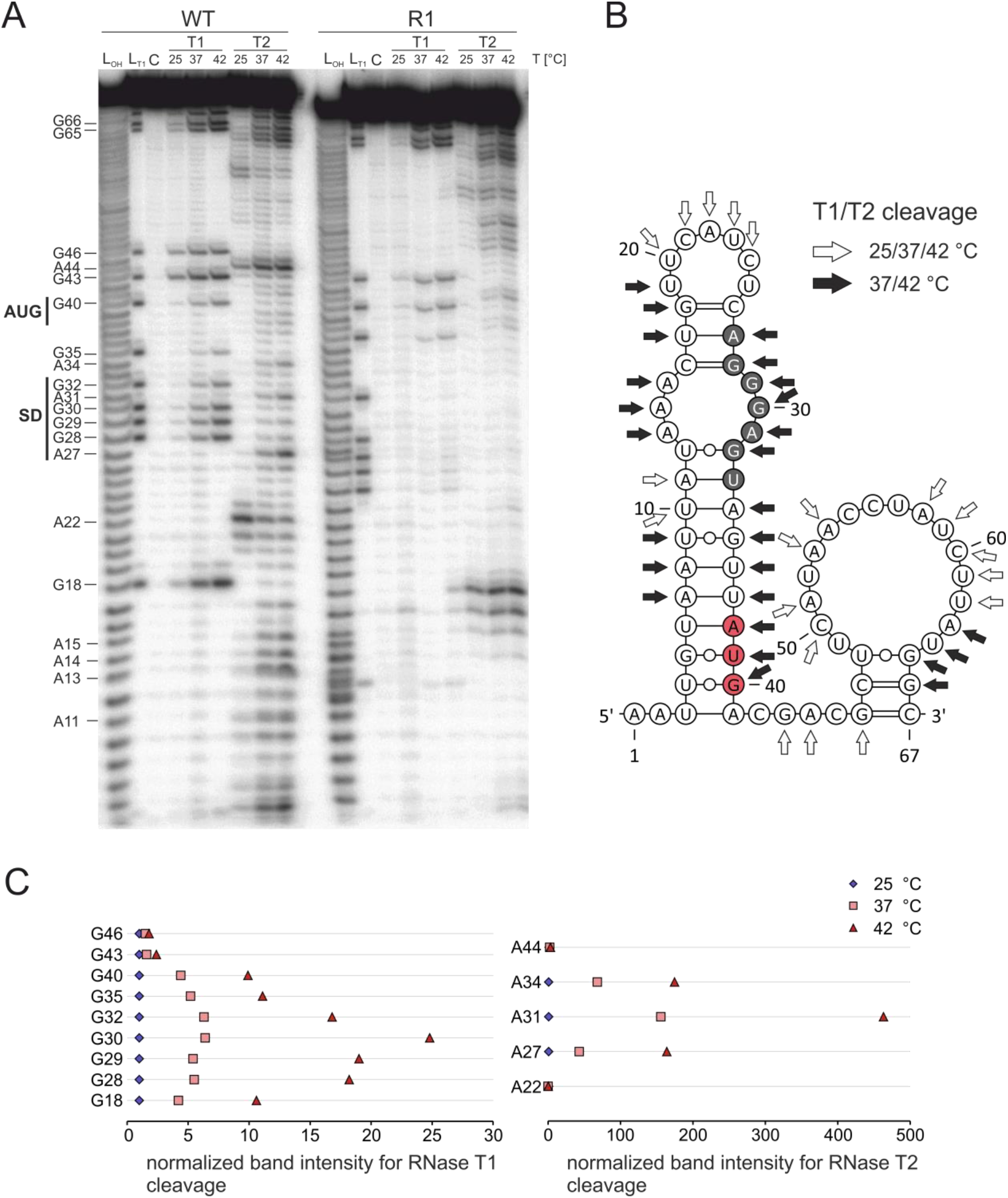
Temperature-dependent melting of the SD sequence and Start codon-containing stem-loop structure of the *yopN* RNAT. (A) Enzymatic structure probing of the *yopN* RNAT (WT) and the stable variant (R1) at 25, 37 and 42 °C. Radioactively labelled RNA was treated with single-strand specific RNases T1 (0.016 U) and T2 (0.025 U) at the different temperatures. L_OH_: alkaline Ladder; L_T1_: T1 treated RNA at 37 °C; C: water control. (B) PARS-derived RNA secondary structure of the *yopN* RNAT [37]. White arrows indicate T1 and T2 cleavages at all temperatures while black arrows indicate cleavages with increasing temperature. The putative SD region is highlighted in gray and the start codon in red. (C) Quantification of band intensities at 25, 37 and 42 °C of selected guanines and adenines. Pixel counting was performed using AlphaEaseFC software and values were normalized to intensities at 25 °C.

### The *yopN* RNAT controls ribosome binding

Typical RNATs adjust expression of the downstream gene to the ambient temperature by controlling the access of the 30S ribosomal subunit to the SD sequence. To demonstrate this activity for the *yopN* RNAT, the short WT and R1 RNAs were subjected to toeprinting (*in vitro* primer extension inhibition) analysis at 25, 37 and 42 °C. Incubation of each RNAT variant with the 30S subunit at these temperatures was followed by reverse transcription. The presence of a truncated cDNA product (toeprint) is indicative of successful binding of the 30S ribosome to the RNA, which generates a roadblock for the reverse transcriptase. Such a toeprint signal at the appropriate position between nucleotides 12 to 14 downstream of the start codon was observed for the WT RNA at 37 and 42 °C (Fig. 7). Consistent with a stabilized structure, this toeprint signal was absent when the R1 RNA was assayed. Instead, another cDNA product was generated under all conditions even in samples without the 30S subunit. Such ribosome-independent formation of termination products is frequently observed [34,40,44] and indicates the presence of a stable hairpin that interferes with reverse transcriptase activity.

**Fig 7.**
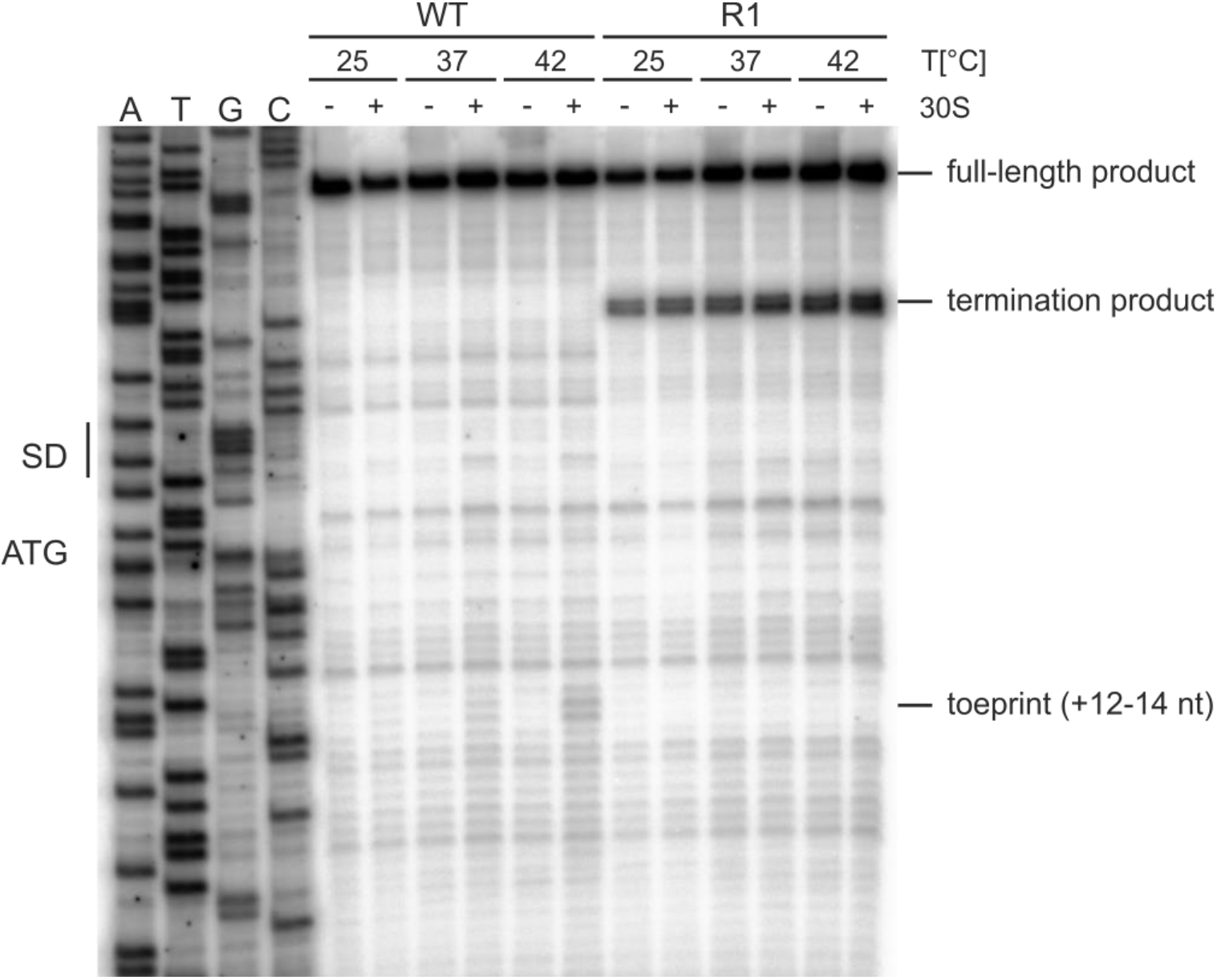
Temperature-dependent binding of the 30S ribosomal subunit to the *yopN* RNAT. Primer extension inhibition of the *yopN* RNAT and the stabile variant R1 was performed at 25, 37 and 42 °C in presence (+) and absence (-) of the 30S ribosomal subunit. A DNA sequencing ladder (ATGC) of the *yopN* RNAT serves as orientation. Full-length transcripts, termination and the characteristic toeprint signals (+12 – 14 nt) after ribosome binding are indicated.

### Unrestricted secretion and inefficient translocation of effector proteins when the RNAT is stabilized

To study the relevance of the *yopN* thermometer on effector protein secretion, we constructed a nonpolar *yopN* deletion mutant as previously described in Bamyaci *et al.* 2018. This Δ*yopN* strain was complemented with plasmids expressing C-terminally Strep-tagged *yopN* either with its natural RNAT or the R1 variant downstream of an arabinose-inducible promoter. Growth of these cultures at 25 °C under non-secretion(+Ca^2+^) or secretion (-Ca^2+^) conditions was indistinguishable from growth of the WT or Δ*yopN* strain carrying the empty vector pGM930 (Fig. 8A). At 37 °C, the previously described temperature-sensitive phenotype of the Δ*yopN* mutant [27] was observed, which is due to the continuous secretion of Yops under both secretion and non-secretion conditions (Fig. 8B). YopN-restricted secretion allowed the WT to grow at 37 °C under non-secretion conditions. The WT-like growth and secretion phenotypes were restored when the Δ*yopN* strain was complemented with the *yopN* expression plasmid carrying the native RNAT. In contrast, uncontrolled permanent secretion and temperature sensitivity of Δ*yopN* could not be rescued when *yopN* was preceded by the closed RNAT variant R1. Western blot analysis taking advantage of the C-terminal Strep-tag of YopN showed that the R1 UTR prevented production of the gatekeeper protein whereas the WT UTR allowed its production specifically at host body temperature (Fig. 8C). Notably, YopN production was further elevated under secretion conditions at low Ca^2+^ concentration.

**Fig 8.**
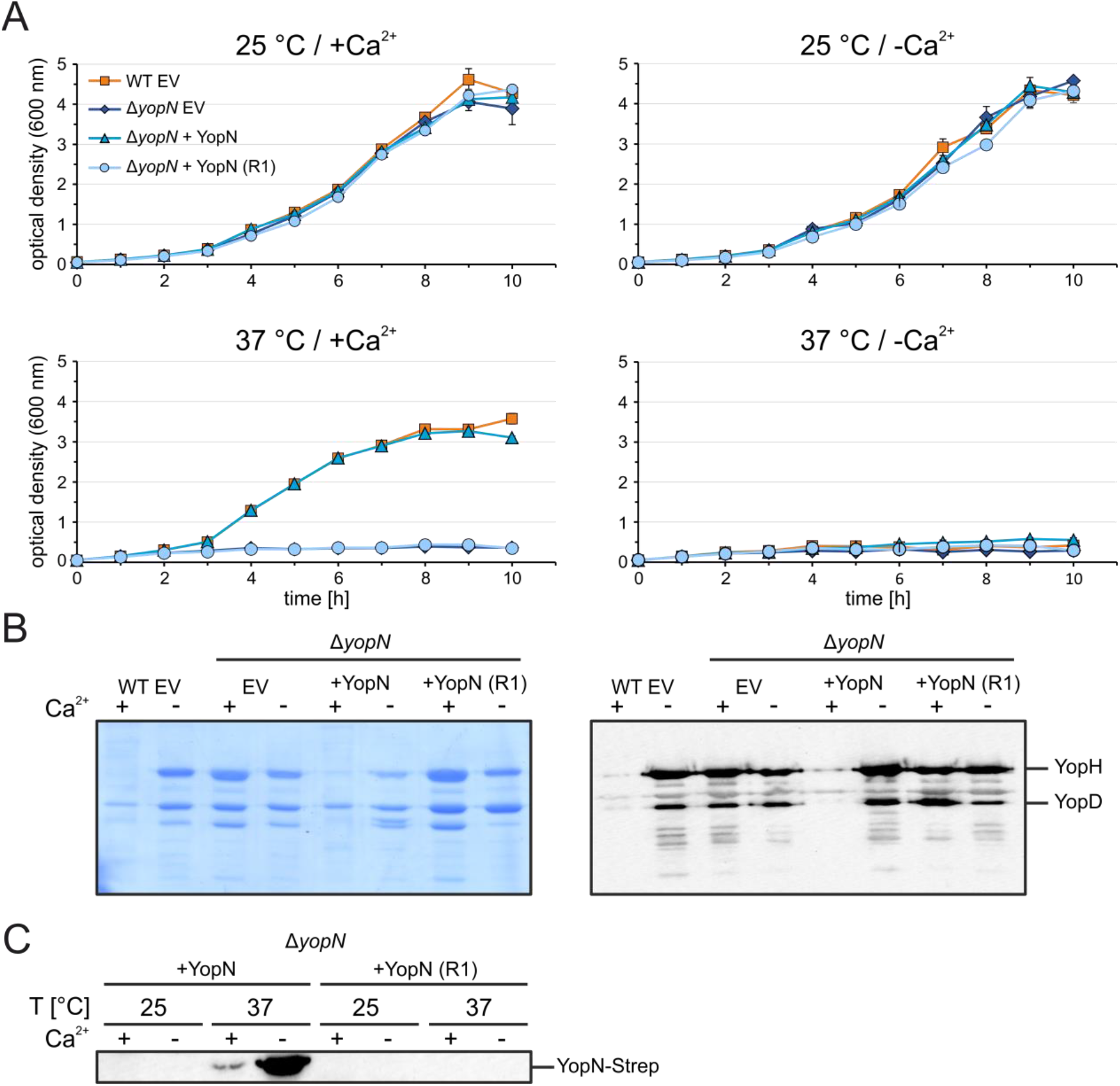
Maintenance of the Δ*yopN* temperature-sensitive phenotype after complementation with the stable RNAT variant R1. (A) Growth experiment of *Y. pseudotuberculosis* YPIII wild type (WT) and Δ*yopN* with the empty vector pGM930 (EV) and Δ*yopN* with vectors containing arabinose-inducible complementation constructs of *yopN* with the wild type RNAT or the stable variant R1. The strains were incubated in secretion-induced (-Ca^2+^) and secretion-noninduced (+Ca^2+^) LB medium at 25 and 37 °C. Growth was measured by optical density at 600 nm. (B) Visualization of secreted effector proteins by SDS-PAGE using Coomassie blue staining and Western blotting using total-anti-Yop serum. Samples were taken after seven hours at 37 °C and secreted proteins were TCA-precipitated from filtered supernatants. (C) For the complementation strains, production of YopN-Strep was visualized by Western blotting using Strep-tag antibody. Samples were taken as described for (B) and protein levels were adjusted to an optical density of 0.5.

Interestingly, YopE translocation assays demonstrated that sustained secretion of Yops in the Δ*yopN* mutant and the mutant complemented with R1 variant did result in enhanced translocation of YopE-TEM into HEp-2 cells (Fig. 9). HEp-2 cells infected with both strains showed almost no blue fluorescence, as a measure of efficient YopE-TEM translocation, compared with wild type and YopN complementation. Concomitantly, HEp-2 cells exhibited a less rounded shape due to the abolished *(ΔyopN)* or reduced (R1) YopE-TEM translocation.

**Fig 9.**
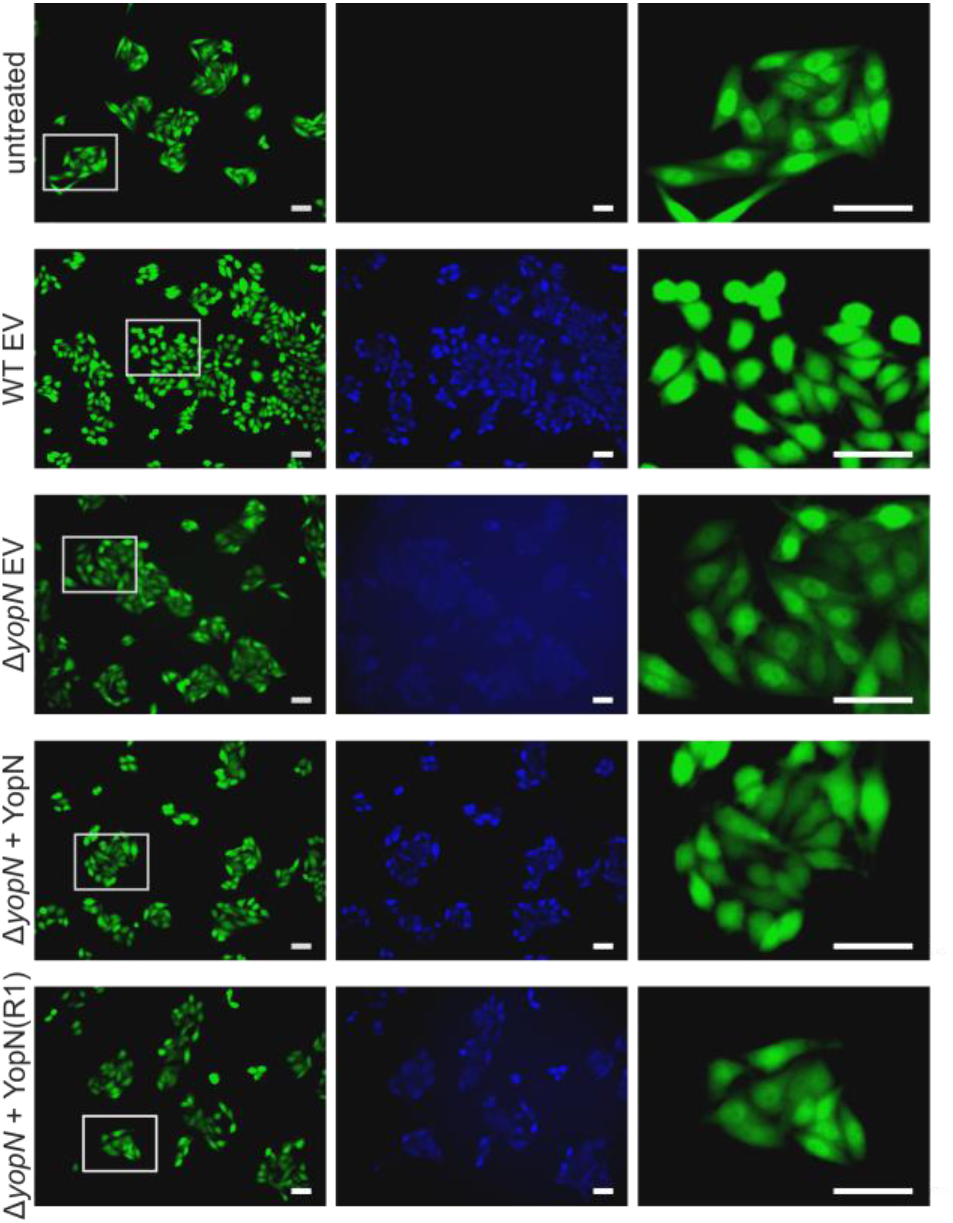
Visualization of YopE translocation into Hep-2 cells. YopE translocation assay of *Y. pseudotuberculosis* YPIII wild type (WT) and Δ*yopN* with the empty vector pGM930 (EV) and Δ*yopN* with vectors containing arabinose-inducible complementation constructs of *yopN* with the wild type RNAT or the stable variant R1. All strains carry plasmid pMK-Bla coding for a *yopE-bla*_TEM_ fusion. HEp-2 cells were infected with bacterial strains at an MOI of 50, labeled with CCF4-AM and analyzed by fluorescence microscopy. Blue fluorescence signals of HEp-2 cells indicate efficient YopE-TEM translocation. In addition, enlarged image sections are shown that highlight the cell morphology. scale bars: 50 μm.

## Discussion

### Multifactorial control of T3SS in *Yersinia*

Many bacterial pathogens use a T3SS as effective syringe-type device to inject effector proteins into host cells in order to manipulate the eukaryotic metabolism to their own benefit. The biosynthesis of the individual components, the correct assembly and dynamic exchange of subunits comes at a considerable cost to the bacterium. Once fully assembled, traffic through the T3SS must be controlled because the unrestricted translocation of effector proteins in the absence of host contact leads to a severe growth arrest (Fig. 8). Growth inhibition might also be due to the loss of essential ions and amino acids through the open T3SS [45–47]. It is therefore not at all surprising that T3S is a tightly regulated process that responds to environmental and host cues with many checks and balances [48].

Since the entire T3SS is encoded on a 70-kbp virulence plasmid, the copy number of this plasmid matters. An immediate strategy of *Yersinia* to boost the expression of T3SS genes under adequate conditions is by increasing the plasmid copy number [49]. Gene dose elevation is followed by a multifaceted regulation of virulence gene expression. A key role in this process plays the ambient temperature because for intestinal pathogens like *Y. pseudotuberculosis*, 37 °C is a reliable indicator of successful invasion of a mammalian host. The expression of more than 300 genes changes at least four-fold between cultures grown at 25 or 37 °C indicating a major temperature-dependent reprogramming of bacterial metabolism and physiology [32]. Temperature-responsive virulence gene expression centers around *lcrF* coding for the primary virulence transcription factor of the Ysc-T3SS/Yop machinery (Fig. 10). Transcription of the *yscW-lcrF* operon at low temperatures outside the host is partially repressed by the global histone-like regulator YmoA. In addition, translation of residual transcripts is inhibited by an RNAT in the intergenic region between *yscW* and *lcrF* [50,51,34]. Three mechanisms contribute to the induction of LcrF levels at 37 °C: (i) a temperature-dependent topology change of the *yscW-lcrF* promoter, (ii) the proteolytic cleavage of YmoA by the Lon and ClpP proteases, and (iii) melting of the *lcrF* RNAT [34,52–54]. To make LcrF-mediated virulence gene induction not solely dependent on the temperature signal, the pathogen has installed additional mechanisms that control the concentration of the transcription factor. Signals indicating host cell contact are integrated via the translocon protein YopD, which – when not secreted – ultimately stimulates *lcrF* mRNA degradation by the degradosome [55].

**Fig 10.**
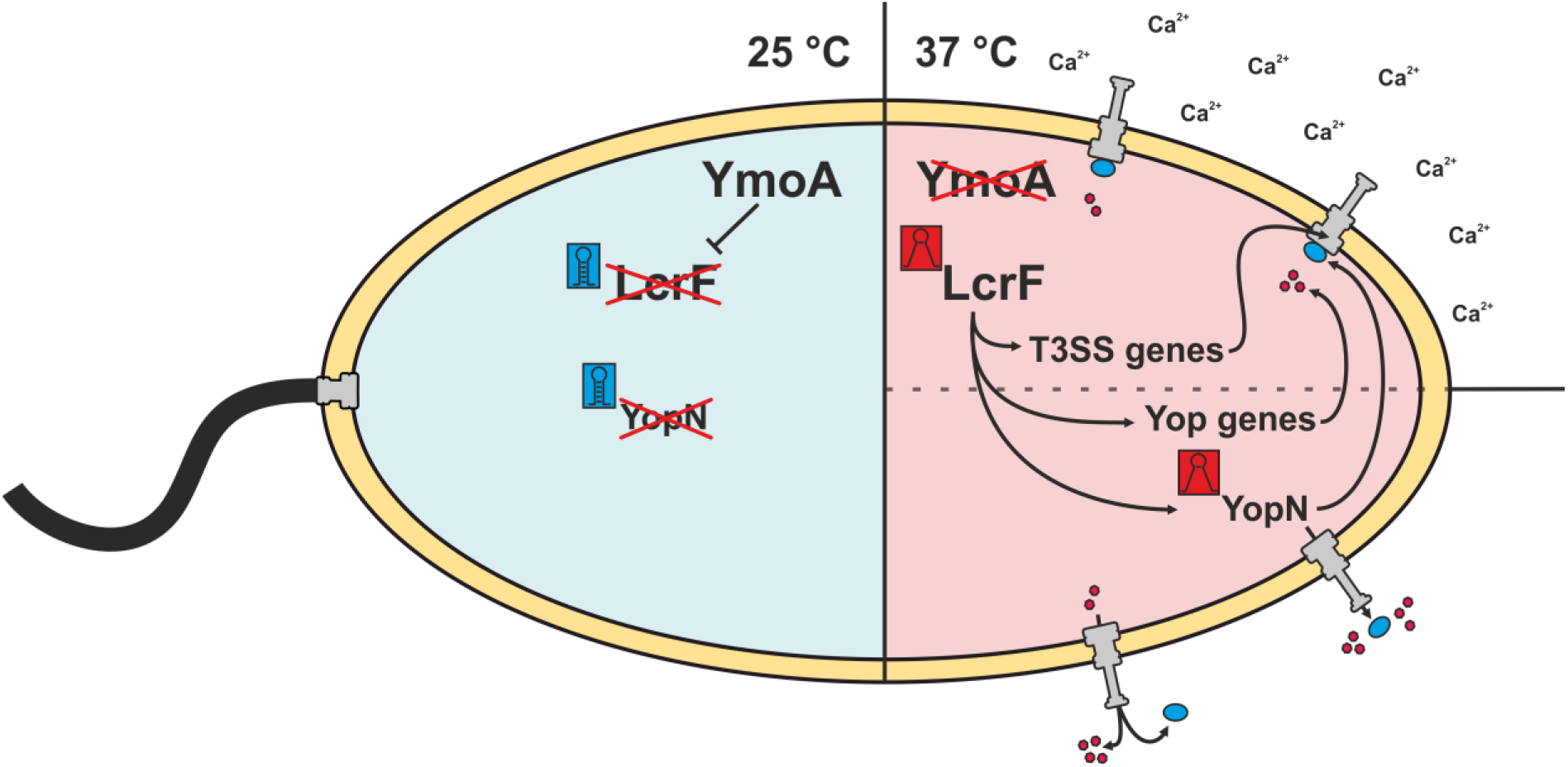
Overview of temperature-dependent regulatory pathways regarding the assembly and functionality of the T3SS. At ambient temperatures (25 °C), *Yersinia* downregulates virulence-associated pathways while the flagellar synthesis is induced [56]. The global regulator YmoA represses the expression of the main virulence regulator *lcrF* at 25 °C [50,52–54]. Besides, RNA thermometers (RNATs) also contribute to repression of specific genes like *lcrF* itself or the secretion regulator *yopN* [34]. At virulence-relevant temperatures (37 °C), YmoA is degraded by proteases which leads to the derepression of *lcrF* [54]. Furthermore, melting of the *lcrF* RNAT increases the expression induction and synthesis of the virulence regulator and thus induce the expression of T3SS genes and effector protein genes (Yop genes) [34]. The transcript of *yopN* posseses an RNAT that additionally induces its expression at 37 °C. In calcium-containing environments, YopN together with TyeA and the chaperone complex SycN/YscB, prevents the secretion of Yops by blocking the T3SS channel in the cytosol. In contrast, the complex dissociates under calcium deficiency allowing first the secretion of YopN and subsequently the secretion of further Yops into the surrounding medium [20,22,23,26,27].

Another important signal in *Yersinia* T3SS gene expression is Ca^2+^, a phenomenon known as low calcium response [57]. The interplay between YopN and its partner proteins (Fig. 1) is responsible for the calcium-controlled secretion of Yops [58]. We observed a combined effect of temperature and low calcium, which mimics the situation in the mammalian host, on *yopN* mRNA (Fig. 2B) and on YopN protein even in the context of a foreign promoter (Fig. 8C) suggesting that both transcription and translation are positively affected by low calcium. Host cell contact-dependent transcriptional regulation of the T3SS and effector proteins involves a number of factors apart from LcrF, including the carbon storage regulator CsrA, the small RNAs CsrB and CsrC and YopD [33,55]. It is possible that this multifactorial network also has an impact on translation efficiency of the *yopN* transcript.

Temperature and host cell contact (simulated by calcium depletion *in vitro)* certainly play a dominant role in T3SS gene expression, but several other environmental parameters influence this process and typically converge on *lcrF* expression. The [2Fe-2S] transcription factor IscR mediates oxygen and iron regulation of the T3SS [59]. It is thought to repress T3SS expression in the intestinal lumen and induce T3SS expression in deeper tissues according to the gradual changes in oxygen tension and iron availability. A role of the CpxAR two-component system [60] and the Rcs phosphorelay system [61], which monitor the bacterial cell envelope integrity, in LcrF induction indicates that various other environmental cues are fed into T3SS regulation. It seems that *Yersinia* has coopted an array of regulatory pathways to sense and integrate the overwhelming signal complexity after transition from the outside to the mammalian gastrointestinal tract in order to precisely adjust the synthesis and activity of the costly and potentially deleterious T3SS to the ambient conditions.

In contrast to upregulation of the T3SS, elevated temperature has an antagonistic effect on flagellum biosynthesis. Motility has been proposed to be a liability in the mammalian host, in part because the flagellum is a potent inducer of innate immunity [62]. Coinciding with virulence gene induction in *Y. enterocolitica* at 37 °C, the transcription of late flagella genes is immediately arrested [56]. Inverse regulation of virulence and motility in *Y. pseudotuberculosis* involves the LysR-type transcription factor RovM [63].

### A novel RNAT at a critical checkpoint of T3SS

The contribution of our study to understanding the intricate control of *Yersinia* virulence lies in the discovery of yet another layer of temperature regulation of the *Yersinia* T3SS. Unlike LcrF-mediated control, this mechanism does not concern the expression of the entire secretion machinery but addresses a highly specific process, the gating of the secretion channel. We identified essentially the same RNAT in both biological isoforms of the *yopN* 5’-UTR, of which the short one (37 nts) is much more abundant than the longer one (102 nts). A functional RNAT in both a long and a short UTR also exists upstream of *Y. pseudouberculosis cnfY* [40]. Roughly equal activity of the RNAT in the short and long *yopN* transcripts can be explained by an identical arrangement of the structure that sequesters the translation initiation region.

The architecture of the short *yopN* RNAT is rather simple and composed of only a single hairpin (Fig. 2B). This makes it one of the shortest known natural RNATs. Many other RNATs contain several hairpins upstream of the decisive thermolabile structure. The adjoining, often more stable hairpins are believed to aid in proper folding of the weaker regulatory hairpin [35]. RNAT-containing 5’-UTRs with a length below 50 nts are exceptional. One example of such a simple helix with an internal asymmetric loop was found upstream of *Synechocystis hsp17* [64,65]. Sequence-wise, the *Yersinia yopN* thermometer is unique and bears no resemblance to RNATs in *Yersinia* or other bacteria. This is contrasted by the *lcrF* RNAT, which belongs to the fourU thermometer family, in which the SD sequence is paired by four uridines [34,66]. Structure-wise, the *yopN* thermometer follows the most common principle. The SD region folds back onto an upstream region that has already been transcribed and is waiting for interaction. While this might have considerable advantages in co-transcriptional RNA folding and instantaneous repression of translation, there are exceptions to this rule. For example, the SD sequence of the *Neisseria meningitidis fHbp* transcript coding for the factor H binding protein folds onto a downstream sequence in the coding region [67].

The necessary temperature-sensitivity in the *yopN* thermosensor is built in by three mismatches opposite the SD sequence. Partial stabilization of this internal loop by a central CG pair was not sufficient to eliminate temperature responsiveness (R2 in Fig. 3). Closing the entire loop by three nucleotides complementary to the SD sequence impaired melting and translation of the *yopN* mRNA at host body temperature (R1 in Fig. 3 to 7) and resulted in massive leakage of Yops accompanied by growth cessation (Fig. 8). In line with other reports [34,40,68–71], these results show how the manipulation of just a few nucleotides in the non-coding region of a protein-coding transcript can dramatically influence expression and the corresponding biological outcome. The results also reinforce the concept that RNAT function depends on a delicate balance of stabilizing and destabilizing elements that render the RNA structure responsive to temperature fluctuations within a narrow and physiologically permissive temperature range. Previous NMR studies provided in-depth insights into the critical contribution of destabilizing internal bulges and loops and non-Watson-Crick-type base pairs to the functional design of various heat shock and virulence thermometers [65,72–74].

The position of the *yopN* RNAT at the beginning of a long mRNA suggests that the six downstream genes of the operon are unaffected by this translational control element. RNAT-mediated differential regulation of the first or second gene of a bistronic operon has been reported for the *Salmonella groESL* operon and *Yersinia yscW-lcrF* operon, respectively [34,75]. RNATs or other riboregulators, such as riboswitches or small regulatory RNAs, provide a simple means to differentially control individual genes in complex operon arrangements [76].

Why is it important to have *yopN* expression under such strict regulation? The conservation of the 5’-UTR sequence in various *Yersinia* species strongly supports its functional relevance. Otherwise, this sequence outside the coding region would be free to evolve. It is conceivable that the additional checkpoint imposed by the RNAT slightly delays the synthesis of YopN in comparison to other T3SS proteins that are all under the regulatory umbrella of LcrF. In the sequential assembly of the T3SS it might be counterproductive to have the gatekeeper present when the T3SS is not ready yet. Despite its limited size, YopN has at least four protein interaction partners, namely TyeA, SycN and YscB (Fig. 1), and YopI, a component of the inner rod of the T3SS [77]. The appropriate order of intermolecular interactions between these proteins might require the precise adjustment of the cellular concentration of the individual components. Another possibility is that YopN has presently undisclosed functions that need to be kept in check when the conditions are not appropriate. Recent studies suggest that YopN has functions beyond its role as gatekeeper of the T3SS. The centrally located coiled-coil domains of YopN encompassing amino acids 65 – 100 provides a virulence-related function. The region contributes to the translocation of YopE and YopH, and is required for systemic infection in mice [29,30].

Our data suggests that YopN is an active rather than passive gatekeeper that exerts a directional influence on Yop secretion. The absence of YopN triggered an uncontrolled premature burst of Yops into the medium, which prevented efficient targeting into host cells. Reduced YopE-TEM translocation by the Δ*yopN* strain with or without the complemented R1 variant was associated with less rounded epithelial cells compared to infections with the WT strain. Delayed rounding of HeLa cells infected with the Δ*yopN* strain has previously been observed [78]. In combination with the finding that YopN is involved in the secretion of YopH and YopE [29,30], these results suggest that YopN regulates the secretion hierarchy to accomplish an ordered translocation of Yops into host cells. Precise timing of the initiation and termination of T3S is critical to efficiently interfere with deleterious immune cell function such as phagocytosis after secretion. While several facets of the biological role of YopN remain unexplored, the cumulative results of the present and other studies suggest functions of this versatile protein that are worth further exploration. Since the incorrect intracellular concentration of YopN causes severe phenotypes, any strategy interfering with the provision of this protein might be suited to combat the pathogenic outcome of *Yersinia* infections

## Material and methods

### Strains, plasmids and oligonucleotides

Bacterial strains and plasmids used in this study are listed in Table S1 and oligonucleotides are listed in Table S2. Bacteria were grown in LB medium and incubated on LB plates at the indicated temperatures. For plasmid-carrying bacteria, the following final antibiotic concentrations were applied: ampicillin (100 μg/mL), kanamycin (50 μg/mL), chloramphenicol (20 μg/mL) and gentamicin (10 μg/mL).

### Plasmid construction

The plasmids pBAD2-*bgaB* and pBAD-*gfp* were served as backbone for reporter gene constructs (Table S1). The short 5’-UTR of *yopN* plus 30 nucleotides of the coding region were amplified with primers yopN_short_fw and yopN_rv and cloned into NheI and EcoRI digested plasmids to obtain translational RNAT:*bgaB* and RNAT:*gfp* fusion constructs (pBO6202 and pBO6207). The long 5’-UTR of *yopN* plus 30 nucleotides of the coding region was cloned into pBAD2-*bgaB* using primers yopN_long_fw and yopN_rv according to the cloning strategy described above (pBO6203).

Plasmids containing a T7 promotor upstream of the short 5’-UTR of *yopN* were constructed for *in vitro* synthesis for enzymatic structure probing and primer extension inhibition. For this purpose, the short 5’-UTR of *yopN* was amplified with primers yopN_rnf_toe_fw and yopN_rnf_rv (pBO6247), and yopN_rnf_toe_fw and yopN_toe_rv (pBO6275) and cloned into SmaI digested pK18.

Specific primers were used to introduce mutations into plasmids by site-directed mutagenesis to obtain the following RNAT variants: R1 (AAA13-15UCC) using yopN_R1_fw and yopN_R1_rv (pBO6256, pBO6269, pBO6297, pBO6270, pBO7273 and pBO6297), R2 (A14C) using yopN_R2_fw and yopN_R2_rv (pBO6257), R3 (UG4-5CA) using yopN_R3_fw and yopN_R3_rv (pBO6258), D1 (C16A) using yopN_D1_fw and yopN_D1_rv (pBO6216) and D2 (CG16,18AA) using yopN_D2_fw and yopN_D2_rv (pBO6255 and pBO6268).

### Generation and complementation of Δ*yopN*

A nonpolar *yopN* deletion strain was constructed as described in Bamyaci *et al*. 2018. In this mutant, a region corresponding to amino acids 9 – 272 of YopN is deleted to maintain the function of the downstream located *tyeA* gene. For this purpose, a 512 bp long 5’-flank and a 446 bp 3’-flank were amplified with primers yopN_5’_fw, yopN_5’_rv, yopN_3’_fw and yopN_3’_rv and recombined by SOE-PCR [79]. The generated deletion fragment was cloned in the suicide plasmid pDM4 using the SacI restriction site [80]. To obtain *yopN* deletion mutants, conjugation with *E. coli* S17-1 *λ-pir* (donor strain) and *Y. pseudotuberculosis* YPIII (recipient strain) was followed by sucrose selection. Putative Δ*yopN* clones were tested by PCR with internal and external primer combinations and confirmed by DNA sequencing (Fig. S2).

For complementation of Δ*yopN,* a construct of the short 5’-UTR, the coding region and a C-terminal Strep-II tag was amplified with the primers yopN_comp_strep_fw and yopN_comp_strep_rv, and ligated into NcoI and PstI digested pGM930 (pBO7423). Using this plasmid, expression of *yopN* is induced by the addition of 0.05 % (w/v) L-arabinose [81]. To obtain a functional complementation with a closed *yopN* RNAT, the R1 variant (AAA13-15UCC) was generated with primers yopN_R1_fw and yopN_R1_rv by site directed mutagenesis (pBO7440).

### RNA isolation

Cell pellets from 4 mL culture samples were resuspended in 250 μL TE buffer (1 mM EDTA, 10 mM Tris, pH 7.5) and 12.5 μL SDS (10% w/v). Then, 450 μL of phenol was added and the samples were incubated at 60 °C for 10 min. After 1 h on ice and centrifugation (1 h, 13000 rpm, 4 °C), 450 μL phenol and 43 μL sodium acetate (3 M, pH 5.5) were added to the aqueous phase followed by centrifugation (5 min, 13000 rpm, 4 °C). The aqueous phase was then mixed with 450 μL chloroform and centrifuged (5 min, 13000 rpm, 4 °C). This step was repeated. After ethanol precipitation and drying for 20 min at 30 °C, the RNA pellets were resuspended in 40 μL sterile water (Carl Roth GmbH, Karlsruhe, Germany).

### Quantitative real-time PCR (qRT-PCR)

For the analysis of relative transcript levels of the short and long 5’-UTR of *yopN,* samples of *Y. pseudotuberculois* YPIII cells for RNA isolation were taken during the early exponential phase at an optical densitiy (OD_600_) of 0.5 under non-secretion (2.5 mM CaCl_2_) and secretion (20 mM MgCl_2_, 20 mM sodium oxalate) conditions in LB medium at 25 and 37 °C. Transcript levels were determined from three independent cultures measured in duplicate. RNA was isolated as described above. For further details on how the qRT-PCR was performed, see Twittenhoff *et al.* 2020b. For the calculation of relative *yopN* transcript levels, the primer efficiency corrected method was used [82]. The non-thermoregulated reference genes *nuoB* and *gyrB* served for normalization [32]. Primer efficiencies were calculated by the CFX Maestro software (*nuoB*: 100.8 %, *gyrB:* 101.3 %, *yopNshort:* 95.3 %, *yopNlong:* 102.8 %).

### Northern blot analysis

Northern blot experiments were carried out as described in [83]. pBAD2-*gfp* served as a DNA template for the amplification of a 286 bp fragment possessing a T7 RNA promotor. Based on the described fragment, an *in vitro* produced and DIG-labeled *gfp* RNA probe (Roche, Mannheim, Germany) was prepared for the detection of *gfp* transcripts.

### Reporter gene assays

*Y. pseudotuberculosis* YPIII cells carrying plasmids of *bgaB* fusion constructs and mutated variants were grown in LB with ampicillin to an OD_600_ of 0.5 at 25 °C. Subsequently, the transcription was induced with 0.1 % (w/v) L-arabinose and the cultures were split to flasks at 25 °C and prewarmed flasks at 37 °C and incubated for further 30 minutes. Samples containing the *bgaB* fusions were then taken for the β-galactosidase assay and Western blot analysis. The mean activities of Miller Units and the mean standard deviations were calculated from nine biological replicates. The β-galactosidase activity was measured as described in [84].

For cells carrying plasmids with *gfp* fusion constructs, the growth and induction was performed as described for *bgaB*. A total of 4 mL of cell suspension were used for Northern blot analysis and 2 mL of cell suspension were used for Western blot analysis.

### Western blot analysis

For the preparation of crude protein extracts, cell pellets of *Y. pseudotuberculosis* YPIII and Δ*yopN* were resuspended in 1 x SDS sample buffer (2% (w/v) SDS, 12.5 mM EDTA, 1% (v/v) β-mercaptoethanol, 10% (v/v) glycerol, 0.02% (w/v) bromophenol blue, 50 mM Tris, pH 6.8) and protein amounts were adjusted by OD_600_ (50 μL per OD_600_ of 1). Resuspended protein samples were heated at 95 °C for 10 min, centrifuged (5 min, 13000 rpm) and loaded to a 12.5% SDS polyacrylamide gel. After SDS-PAGE, proteins were blotted onto a nitrocellulose membrane (Hybond-C Extra, GE Healthcare, Munich, Germany) Ponceau S staining of the blotted membrane was performed to assure equal amounts of proteins. For the detection of BgaB-His, a penta-His HRP conjugate was used (1:4000, QIAGEN GmbH, Hilden, Germany). The detection of GFP was performed with the primary antibody anti-GFP (1:10000, Thermo Scientific, Waltham, USA) followed by the secondary antibody goat anti-rabbit HRP conjugate (1:4000, Bio Rad, Munich, Germany). In the case of YopN-Strep, the protein was detected using the strep-tactin HRP conjugate (IBA Lifesciences, Göttingen, Germany) as specified by the manufacturer. Furthermore, for the visualization of secreted Yops, a total Yop antiserum [39] was used (1:20000) followed by the secondary antibody goat anti-rabbit HRP conjugate (1:4000, Bio Rad, Munich, Germany). Protein signals on the membranes were detected with Immobilon Forte Western HRP substrate (Merck, Darmstadt, Germany) and the ChemiImager Ready (Alpha Innotec,San Leandro USA).

### Enzymatic structure probing

To enzymatically probe RNA structures at different temperatures, RNA was first synthesized by *in vitro* run-off transcription using T7 RNA polymerase (Thermo Scientific, Waltham, USA) and the EcoRV-linearized plasmids as described in Righetti *et al.,* 2016. For structure probing, the plasmids pBO6247 and pBO6270 were used to synthesize RNA consisting of the short 5’-UTR of *yopN* plus 30 nt of the coding region and the variant R1. Purified and desphosphorylated RNA was labeled with [^32^P] at the 5’-end as described in [85]. According to [66], limited digestion of radiolabeled RNA was performed with the ribonuclease T1 (0.005 U) (Thermo Scientific, Waltham, USA) and T2 (0.074 U) (MoBiTec, Göttingen, Germany) in 5 x TN buffer (500 mM NaCl, 100 mM Tris acetate, pH 7) at 25, 37 and 42 °C. Furthermore, an alkaline hydrolysis ladder [85] and a T1 ladder was prepared. For the T1 ladder, 30000 cpm of labeled RNA was heated with 1 μL sequencing buffer (provided with RNase T1) to 90 °C and then incubated with the nuclease at 37 °C for additional 5 min.

### Primer extension inhibition assay (toeprinting)

*in vitro* transcribed RNA produced with the plasmids pBO6265 and pBO6273 (short 5’-UTR of *yopN* plus 60 nt of the coding region and variant R1), 5’-[^32^P]-labeled reverse primer yopN_short_toe_rv, 30S ribosomal subunits and tRNA^fMet^ (Sigma-Aldrich, St. Louis, USA) were used for the toeprinting assay as described in [86]. First, the annealing mix consisting of 0.08 pmol RNA and 0.16 pmol radiolabeled primer were mixed with 1 x VD-Mg^2+^ (60 mM NH4Cl, 6 mM β-mercaptoethanol, 10 mM Tris/HCl, pH 7.4) and incubated for 3 min at 80 °C. For binding of 30S ribosomal subunits, the annealing mix was mixed with 16 ρmol tRNA^fMet^, 0.16 ρmol radiolabeled primer and 6 ρmol 30S ribosomal subunit in Watanabe buffer (60 mM HEPES/KOH, 10.5 mM Mg(CH_3_COO)_2_, 690 mM NH_4_COO, 12 mM β-mercaptoethanol, 10 mM spermidine, 0.25 mM spermine) and incubated for 10 min at 25, 37 and 42 °C. Subsequently, a M-MLV mix (1 x VD+Mg^2+^ (10 mM Mg(CH_3_COO)_2_, 6 μg BSA, 4 mM dNTPs, 800 U MMLV reverse transcriptase (Thermo Scientific, Waltham, USA)) was added to initiate the cDNA synthesis for 10 min at 37 °C. After the addition of formamide loading dye, the reaction was stopped and the samples were separated on an 8% polyacrylamide gel. A sequencing ladder was produced with the Thermo Sequenase cycle sequencing Kit (Thermo Scientific, Waltham, USA), the template pBO6265 and the radiolabeled primer yopN_short_toe_rv.

### Growth experiments and Yop secretion

*Y. pseudotuberculosis* YPIII and Δ*yopN* carrying complementation constructs (pGM930) were grown under non-secretion (2.5 mM CaCl_2_) and secretion conditions (20 mM MgCl_2_, 20 mM sodium oxalate) in LB containing 0.05 % (w/v) L-arabinose and ampicillin for 10 h at 25 and 37 °C h. After 7 h, a total of 9 mL was collected, adjusted to an OD_600_ of 0.8 and centrifuged (10 min, 4000 rpm, 4 °C). To the sterile filtered supernatant, 1 mL of 100% (w/v) TCA was added to precipitate secreted proteins overnight at 4 °C followed by a centrifugation (20 min, 13000 rpm, 4 °C). Protein pellets were first resuspended in 0.5 mL 2% SDS solution and then mixed with 1.5 mL of ice-cold 100% acetone. The samples were incubated for 30 min at −20 °C. After washing twice with 0.5 mL acetone, pellets were dried for 15 min at 25 °C. Finally, pellets were resuspended in 25 μL of 2 x SDS sample buffer (4% (w/v) SDS, 25 mM EDTA, 2% (v/v) β-mercaptoethanol, 20% (v/v) glycerol, 0.04% (w/v) bromophenol blue, 100 mM Tris, pH 6.8) and 5 μL were loaded to an 12.5 % SDS acrylamide gel to separate secreted *Yersinia* effector proteins (Yops).

### YopE translocation assay

The YopE translocation assay was performed to examine the YopE secretion ability of *Y. pseudotuberculosis* strains carrying complementation constructs (pGM930) and pMK-Bla (YopE-TEM, [87] using LiveBLAzer-FRET B/G Loading Kit (Life Technologies, Carlsbad, USA). Strains were pregrown in LB containing 0.1 % (w/v) L-arabinose, 1 mM CaCl_2_, ampicillin and kanamycin for 2 h at 25 °C. The bacteria were then shifted to 37 °C, incubated for additional 2 h, washed and adjusted in PBS buffer to an OD_600_ of 1. HEp-2 cells (1.7×10^4^) seeded in μ-Slides (8-well, Ibidi, Gräfelfing, Germany) were infected with *Y. pseudotuberculosis* strains at MOI of 50 and centrifuged for 5 min (400 g, RT). This was followed by incubation for 60 min at 37 °C and three washing steps of the cells with PBS buffer. Then, 200 μL of infection buffer (RPMI, 20 mM HEPES, 0.4% BSA) containing gentamicin (25 μg/mL) were added and HEp-2 cells were stained with CCF4-AM loading dye according to the manufacturer’s protocol. YopE-TEM translocation was visualized using a fluorescence microscope (BZ-9000 Fluorescence Microscope, Keyence, Osaka, Japan).

## Supporting information

**S1 Table. Bacterial strains and plasmids used in this study.** (DOCX)

**S2 Table. Oligonucleotides used in this study.** (DOCX)

**S1 Fig. Sequence alignment of the *yopN* 5’-UTR of different *Yersinia* species.** Sequence comparison of the *yopN* 5’-UTRs and 30 nucleotides of the coding region between *Y. pseudotuberculosis*, *Y. pestis* and *Y. enterocolitica*. Bold black nucleotides indicate the putative SD region and start codon while red nucleotides indicate sequence variations between *Yersinia* species. l: long transcript; s: short transcript. (TIF)

**S2 Fig. Confirmation of the Δ*yopN* mutant by PCR.** (A) Altered DNA fragment sizes due to *yopN* deletion using primers that bind within *yopN* (I:internal primers) and up- and downstream of the gene locus. Due to *yopN* deletion, no DNA fragment is produced with internal primers compared to the wild type (WT). L: DNA ladder; e: external primers; i: internal primers. (B) Schematic overview of primer localization. (TIF)

**S1 References. References for supporting information.** (DOCX)

## Acknowledgments

We thank Johanna Roßmanith for advice in the initial stage of this project, the RNA group for continuous discussions and Alexander Kraus for critical reading of an earlier version of this manuscript.

## Author contributions

**Conceptualization:** Stephan Pienkoß, Petra Dersch, Franz Narberhaus

**Formal analysis:** Stephan Pienkoß

**Funding acquisition:** Franz Narberhaus

**Investigation:** Stephan Pienkoß, Soheila Javadi, Paweena Chaoprasid, Thomas Nolte

**Methodology:** Stephan Pienkoß, Soheila Javadi, Paweena Chaoprasid

**Project administration:** Franz Narberhaus

**Resources:** Petra Dersch, Franz Narberhaus

**Supervision:** Petra Dersch, Franz Narberhaus

**Validation:** Stephan Pienkoß

**Visualization:** Stephan Pienkoß

**Writing - original draft:** Stephan Pienkoß

**Writing - review & editing:** Paweena Chaoprasid, Alexander Kraus, Petra Dersch, Franz Narberhaus

